# The impact of non-cardiomyocyte *MYBPC3* expression on the development of hypertrophic cardiomyopathy

**DOI:** 10.64898/2026.04.20.718297

**Authors:** Nicolas G. Clavere, Jenny H. Kim, Katie P. Letcher, Sejal T. Molakaseema, Kavisha Silva, Soumojit Pal, Jason R. Becker

**Author notes:** Corresponding author: Jason R. Becker, MD, 200 Lothrop, BST E1258, Pittsburgh, PA 15213.

## Abstract

**Introduction:** Hypertrophic Cardiomyopathy (HCM) is a disease defined by the development of left ventricle hypertrophy. One of the most commonly mutated genes in HCM is cardiac myosin binding protein C (*MYBPC3*). MYBPC3 protein localizes to the cardiomyocyte sarcomere, but studies have reported detection of both *MYBPC3* RNA and protein in non-cardiomyocyte cell populations. Therefore, it was unclear if *MYBPC3* expression in non-cardiomyocyte cell populations altered the development of cardiomyopathy caused by MYBPC3 protein deficiency.

**Methods:** We utilized genetically modified murine models with germline deletion of *Mybpc3* exons 3 to 5 (*Mybpc3*^−/−^) or cardiomyocyte specific deletion of *Mybpc3* exons 3 to 5 (*Mybpc3*^fl/fl^; Myh6-Cre). Gene expression was assessed using quantitative RT-PCR. Whole tissue protein levels were assessed using immunoblots. Immunohistochemistry and proximity ligation assays were performed to evaluate in situ protein expression. Echocardiography was utilized to measure left ventricular structure and function.

**Results:** *Mybpc3* mRNA was detected in multiple organs including the heart, lung and blood from both humans and mice. Utilizing transgenic murine models with germline or cardiomyocyte specific deletion of *Mybpc3* exons 3-5, we discovered that the *Mybpc3* mRNA detected in extracardiac locations originated primarily from cardiomyocytes. Likewise, MYBPC3 protein was identified in myocardial tissue but not in other organs and cardiomyocytes were the only cell population in myocardial tissue that had detectable MYBPC3 protein. Importantly, cardiomyocyte deletion of *Mybpc3* caused similar pathological myocardial remodeling and alterations in left ventricular function compared to germline deletion of *Mybpc3* in all cell populations.

**Conclusions:** Our results show that cardiomyocytes are the primary cell source of *Mybpc3* mRNA detected in extracardiac organs and they are the principal cell type responsible for the cardiomyopathy caused by MYBPC3 protein deficiency. These results suggest that selective targeting of cardiomyocytes should be the most efficient approach to treat cardiomyopathies associated with MYBPC3 deficiency.

## Introduction

Hypertrophic cardiomyopathy (HCM) is characterized by the development of the left ventricle hypertrophy and has a prevalence of at least 1 in 500 in humans.^1^ This disease is often inherited, and the most common genetic causes of this condition are mutations in the sarcomere proteins myosin heavy chain 7 (*MYH7*) and cardiac myosin binding protein C (*MYBPC3*).^2^ Mutations in *MYH7* are typically thought to cause disease through a gain of function mechanism.^3,4^ In contrast, mutations in *MYBPC3* are primarily thought to cause disease through a loss of function haploinsufficiency mechanism.^5–7^

MYBPC3 protein has been shown to be an integral component of the cardiomyocyte sarcomere where it regulates the interaction of myosin and actin filaments to modulate sarcomere contraction.^7–9^ However, *MYBPC3* RNA has been detected in multiple extracardiac tissues such as the blood, lung, adrenal gland, and skeletal muscle.^10,11^ Likewise, non-cardiomyocyte cells such as myocardial fibroblasts and the NIH-3T3 cell line were reported to express MYBPC3 protein.^12^ Likewise, murine models deficient in MYBPC3 protein develop myocardial hypertrophy that is associated with alterations in not only cardiomyocytes but also non-cardiomyocyte cell populations.^13,14^ It was assumed that the changes detected in these non-cardiomyocyte myocardial cell populations were secondary to the pathologic changes in cardiomyocyte growth and function resulting from MYBPC3 protein deficiency. However, it remained unclear whether non-cardiomyocytes expressed MYBPC3 protein and if this non-cardiomyocyte expression altered the development of cardiomyopathy caused by MYBPC3 protein deficiency.

Deciphering the impact of non-cardiomyocyte MYBPC3 protein expression is particularly important since emerging methods to treat MYBPC3 related cardiomyopathies selectively target the cardiomyocyte cell population.^15,16^ In order to address this question, we compared a murine model with germline *Mybpc3* deletion in all cells to a murine model with selective cardiomyocyte *Mybpc3* deletion. We used these in vivo models to identify the primary source of extracardiac *Mybpc3* RNA and to determine if non-cardiomyocyte MYBPC3 protein expression impacts the development and progression of cardiomyopathy resulting from MYBPC3 deficiency.

## Results

### Cardiomyocytes are the primary source of extracardiac *Mybpc3* mRNA

We utilized the Genotype Tissue Expression (GTEx) dataset to evaluate human *MYBPC3* RNA expression in cardiac and extracardiac tissues. We discovered that human *MYBPC3* RNA expression was highest in the heart but was also detected in other organs such as blood and lung (**Figure 1A**). Similar to humans, mice also had *Mybpc3* mRNA expression in extracardiac tissues such as the lungs and blood (**Figure 1B, S1A**). To investigate the source of extracardiac *Mybpc3* mRNA, we utilized a transgenic murine model with germline deletion of exons 3 to 5 of the *Mybpc3* gene in all cell types (*Mybpc3*^−/−^) (**Figure 1C**) and a transgenic murine model that eliminates exons 3 to 5 of the *Mybpc3* gene specifically in cardiomyocytes (*Mybpc3*^fl/fl^; Myh6-Cre) (**Figure 1D**). By selectively measuring *Mybpc3* mRNA that contained exons 3 to 5 from these two transgenic models, we could determine the cardiomyocyte versus non-cardiomyocyte source of *Mybpc3* mRNA. We discovered that left ventricle *Mybpc3* mRNA expression was significantly reduced when cardiomyocyte *Mybpc3* mRNA was eliminated (**Figure 1E**). Likewise, lung and blood *Mybpc3* mRNA was also significantly reduced when cardiomyocyte *Mybpc3* mRNA was eliminated (**Figure 1F-G**). Since our blood samples were obtained by direct LV puncture, we also confirmed that *Mybpc3* mRNA was detected in whole blood obtained directly from the aorta (**Figure S1B**). Overall, these results show that extracardiac *Mybpc3* mRNA is derived primarily from cardiomyocytes.

**Figure 1:**
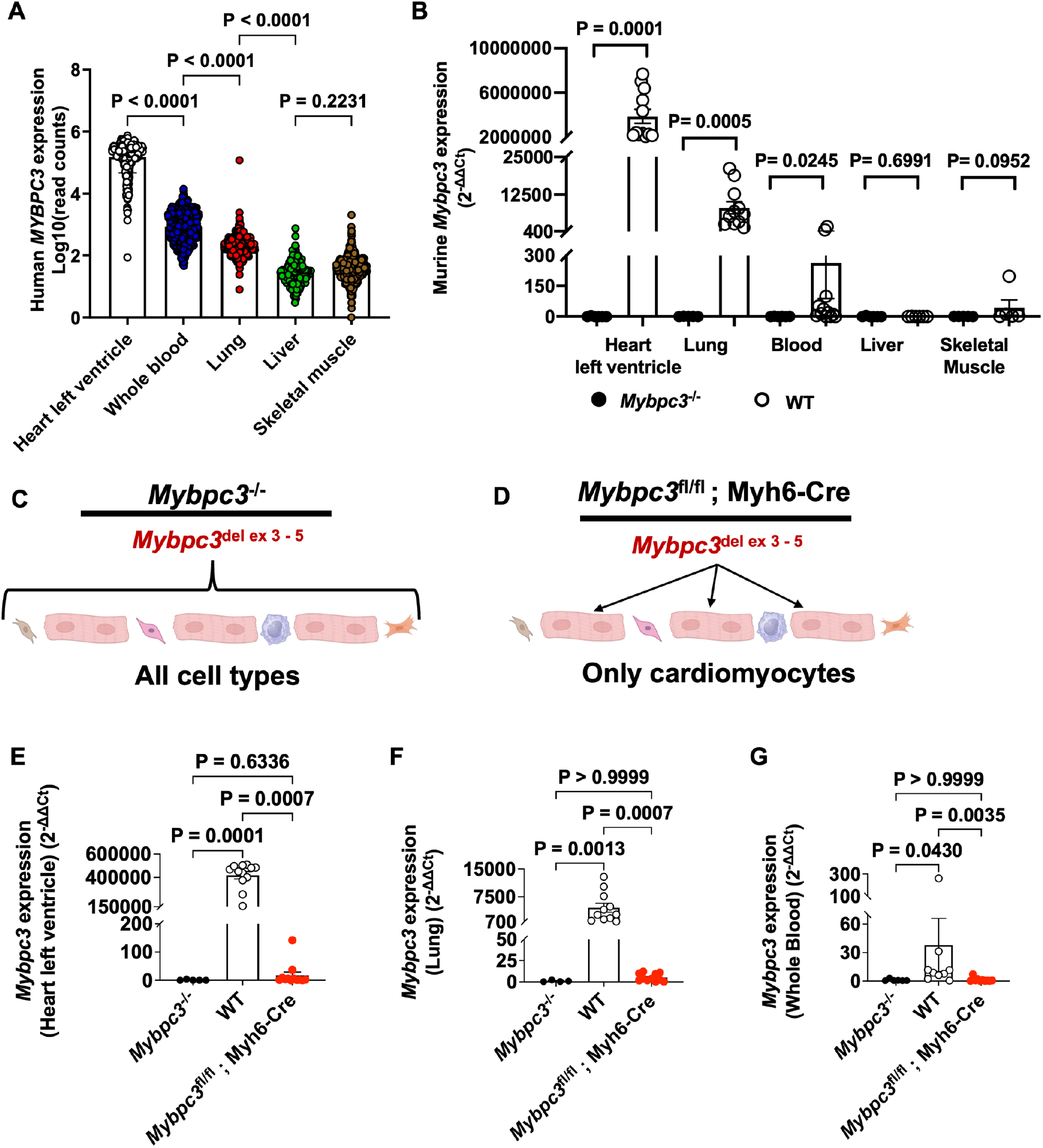
Cardiomyocytes are the primary source of extracardiac *Mybpc3* mRNA. **(A**) Human bulk tissue RNA-sequencing for *MYBPC3* expression from Genotype-Tissue Expression (GTEx) database. *MYBPC3* mRNA expression is displayed as Log10(read counts) for cardiac left ventricle (n=432), whole blood (n=755), lung (n=578), liver (n=226), kidney (n=85), brain (n=255) and skeletal muscle (n=803). **(B)** Quantification of *Mybpc3* expression from *Mybpc3*^−/−^ and WT heart left ventricle (n=6-12), lung (n=5-11), whole blood (n=6-12), liver (n=6) and skeletal muscle (n=5) at postnatal day (P) 90. The *Mybpc3* gene expression was normalized to the housekeeping gene *Rpl32* and the cDNA was created using random hexamer primers. Results are expressed as a fold change relative to *Mybpc3* mRNA expression in the *Mybpc3*^−/−^ mouse tissue. **(C-D)** Schematic of the *Mybpc3* germline knockout (*Mybpc3*^−/−^) and cardiomyocyte specific deletion (*Mybpc3*^fl/fl^; Myh6-Cre). Schematic drawing was performed with BioRender software. **(E)** Quantification of *Mybpc3* expression from *Mybpc3*^−/−^ (n=5), WT (n=12) and *Mybpc3*^fl/fl^; Myh6-Cre (n=12) heart left ventricle at P90. The *Mybpc3* gene expression was normalized to the housekeeping gene *Rpl32*. Results are expressed as a fold change relative to *Mybpc3* mRNA expression in the *Mybpc3*^−/−^ hearts. **(F)** Quantification of *Mybpc3* expression from *Mybpc3*^−/−^ (n=4), WT (n=11) and *Mybpc3*^fl/fl^; Myh6-Cre (n=11) lung at P90. The *Mybpc3* gene expression was normalized to the housekeeping gene expression *Rpl32*. Results are expressed as a fold change relative to *Mybpc3* mRNA expression in the *Mybpc3*^−/−^ lungs. **(G)** Quantification of *Mybpc3* expression from, *Mybpc3*^−/−^ (n=6), WT (n=9) and *Mybpc3*^fl/fl^; Myh6-Cre (n=10) blood at P90. The *Mybpc3* gene expression was normalized to the housekeeping gene expression *Rpl32*. Results are expressed as a fold change relative to *Mybpc3* mRNA expression in the *Mybpc3*^−/−^ blood. All results are shown as mean±SEM. Kruskal-Wallis test with Dunn’s post hoc test for multiple comparisons for was used for **A, E, F** and **G**. Mann Whitney U test was used for **B** to compare WT and *Mybpc3*^−/−^ for heart left ventricle, lung, whole blood, liver and skeletal muscle.

### Extracardiac *Mybpc3* mRNA does not lead to detectable MYBPC3 protein

Since we detected *Mybpc3* mRNA in extracardiac tissues we wanted to determine if this *Mybpc3* mRNA leads to detectable MYBPC3 protein in extracardiac organs. We readily detected MYBPC3 protein in heart left ventricle tissue lysate using two different primary antibodies and two independent imaging methods (**Figure 2A**). In contrast to the left ventricle, we were unable to detect MYBPC3 protein in extracardiac organ lysates such as lung, liver, brain, kidney and skeletal muscle using these same methods (**Figure 2A-C**). In addition, we were unable to detect MYBPC3 protein in whole blood samples (**Figure 2D**). Overall, these results show that extracardiac *Mybpc3* mRNA does not lead to detectable MYBPC3 protein in non-cardiac organs.

**Figure 2:**
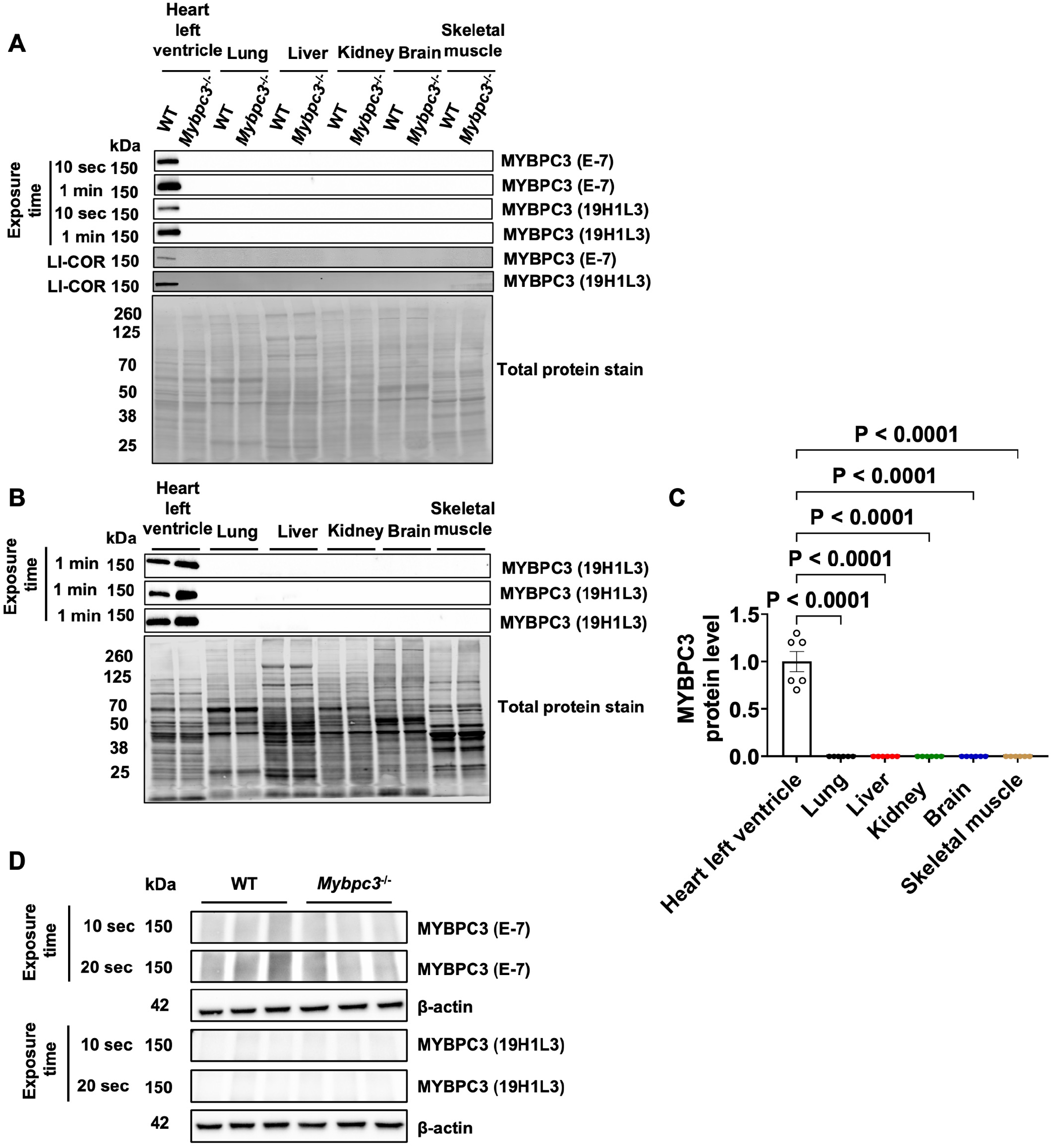
Extracardiac *Mybpc3* mRNA does not lead to detectable MYBPC3 protein. **(A)** Immunoblot images of MYBPC3 protein expression from WT and *Mybpc3*^−/−^ mice in heart left ventricle, lung, liver, kidney, brain and skeletal muscle from mouse tissue at post-natal day 180 (P180). The total protein stain was used as the loading control. The immunoblots used MYBPC3 antibodies 19H1L3 (Invitrogen, 703574) or E-7 (Santa Cruz Biotechnology, sc-137180) and were imaged with ChemiDoc (exposure time detailed) or LI-COR imaging systems. **(B)** Representative immunoblot images of MYBPC3 protein expression from WT mice in heart left ventricle, lung, liver, kidney, brain and skeletal muscle from mouse tissue at P180. The total protein stain was used as the loading control. The immunoblot used MYBPC3 antibody 19H1L3 (Invitrogen, 703574) and was imaged with ChemiDoc (exposure time detailed). **(C)** MYBPC3 protein quantification from WT (n=6) left ventricle, lung, liver, kidney, brain and skeletal muscle mouse tissue at P180. **(D)** Immunoblot images of MYBPC3 protein expression from WT and *Mybpc3*^−/−^ mice in whole blood from aortic puncture using a MYBPC3 antibody 19H1L3 (Invitrogen, 703574) and imaged with ChemiDoc (exposure time detailed). The housekeeping protein β-actin was used as the loading control. All results are shown as mean±SEM. Kruskal-Wallis test with Dunn’s post hoc test for multiple comparisons was used for **C**.

### Non-cardiomyocyte cells of the myocardium do not express MYBPC3 protein

Since myocardial tissue was identified as the primary source of MYBPC3 mRNA and protein, we next wanted to determine if non-cardiomyocyte cells had detectable MYBPC3 protein. We again utilized our transgenic model that eliminated *Mybpc3* exon 3-5 expression in all cells (Mybpc3^−/−^) or specifically in cardiomyocytes (Mybpc3^fl/fl^; Myh6-Cre). First, we used immunohistochemistry and found that cardiomyocyte specific elimination of *Mybpc3* led to no detectable myocardial MYBPC3 protein (**Figure 3A-B**). Next, we performed proximity ligation assays to quantify the in-situ MYBPC3 protein expression in cardiomyocytes and non-cardiomyocyte cell populations. Similar to the IHC experiment, cardiomyocytes had MYBPC3 protein complexes, but non-cardiomyocytes did not (**Figure 3C-D, S2A**). Likewise, human myocardial tissue cardiomyocytes expressed MYBPC3 protein, but non-cardiomyocytes did not have detectable MYBPC3 protein expression (**Figure 3E-F**). To confirm the in situ data, we also performed immunoblots on left ventricle tissue lysate from mice with cardiomyocyte specific deletion of *Mybpc3* and did not detect any residual MYBPC3 protein (**Figure 3G-H, S2B**).

**Figure 3:**
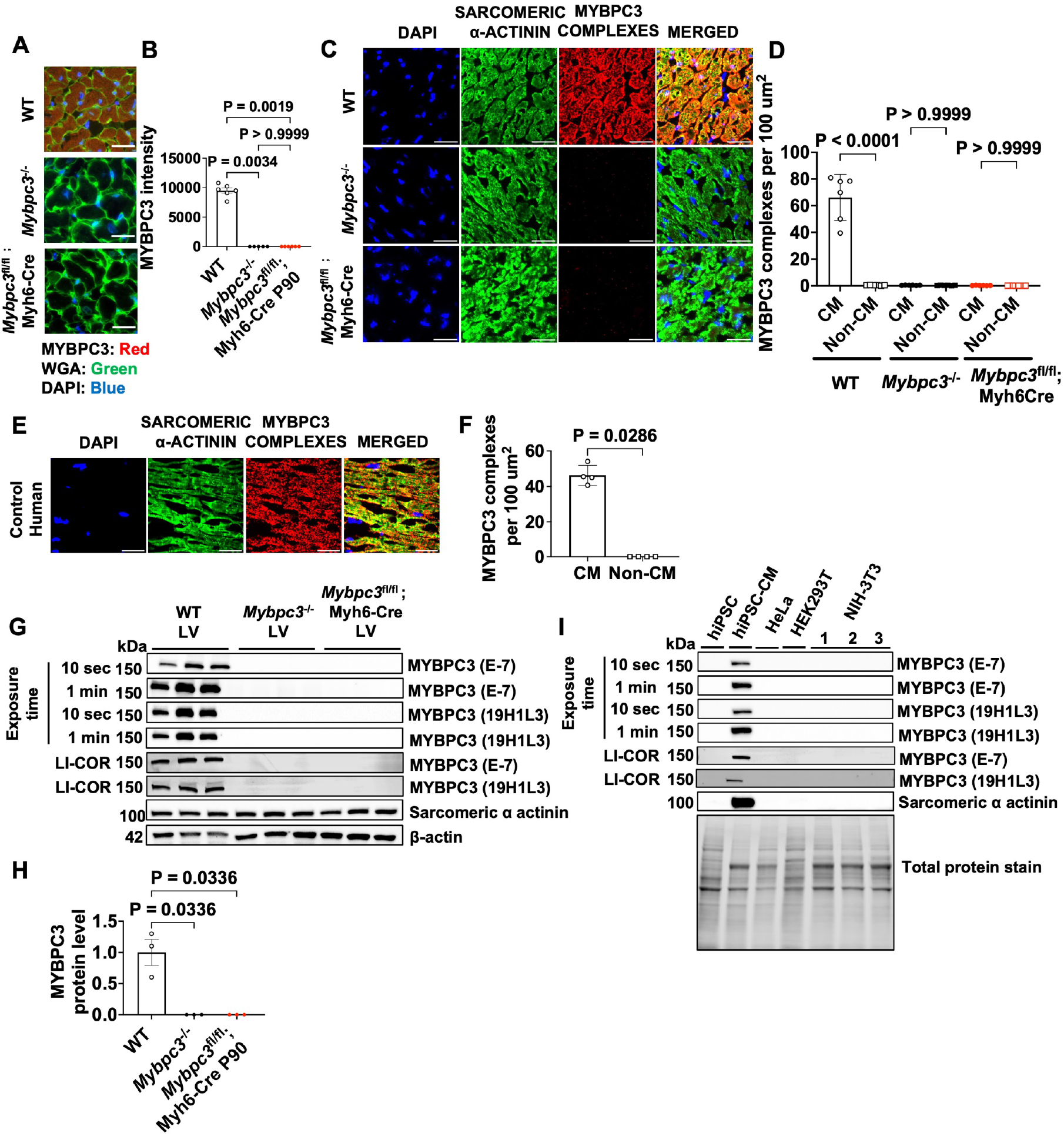
Non-cardiomyocyte cells of the myocardium do not express MYBPC3 protein. **(A)** Representative images of immunofluorescence staining of WT, *Mybpc3*^−/−^ and *Mybpc3*^fl/fl^; Myh6-Cre left ventricle mouse tissue at postnatal day 90 (P90). MYBPC3 - red, wheat germ agglutinin (WGA) - green, 4’6-diamidino-2-phenylindole (DAPI) - blue Scale bars, 25μm. **(B)** MYBPC3 fluorescence intensity quantification from WT (n=6), *Mybpc3*^−/−^ (n=5) and *Mybpc3*^fl/fl^; Myh6-Cre (n=6) at P90. Minimum of 100 cardiomyocytes/sample. **(C)** Representative images of a in situ proximity ligation assay for MYBPC3 (red) counter stained with sarcomeric α actinin (green) and DAPI (blue) in WT, *Mybpc3*^−/−^ and *Mybpc3*^fl/fl^; Myh6-Cre left ventricle tissue at P180. Scale bars, 25μm. **(D)** In situ proximity ligation assay to quantify MYBPC3 protein complexes (per 100μm^2^) in cardiomyocytes *versus* non-cardiomyocytes in WT, *Mybpc3*^−/−^ and *Mybpc3*^fl/fl^; Myh6-Cre (n=6/group) left ventricle tissue at P180. **(E)** Representative images of an in situ proximity ligation assay for MYBPC3 (red) counter stained with sarcomeric α actinin (green) and DAPI (blue) in human control (n=4) left ventricle. Scale bars, 25μm. **(F)** Quantification of MYBPC3 complexes (complexes per 100μm^2^) in cardiomyocytes *versus* non-cardiomyocytes in human control (n=4) left ventricle. **(G)** Immunoblot images of MYBPC3 and Sarcomeric α actinin protein expression from WT, *Mybpc3*^−/−^ and *Mybpc3*^fl/fl^; Myh6-Cre mice left ventricle tissue lysate at P180. The immunoblots used MYBPC3 antibodies 19H1L3 (Invitrogen, 703574) or E-7 (Santa Cruz Biotechnology, sc-137180) and were imaged with ChemiDoc (exposure time detailed) or LI-COR imaging systems. β-actin was used as a loading control. **(H)** MYBPC3 protein quantification from WT, *Mybpc3*^−/−^ and *Mybpc3*^fl/fl^; Myh6-Cre (n=3/group) left ventricle mouse tissue at P180. Quantification was performed using the immunoblot image from antibody 19H1L3 with 1 minute exposure time on ChemiDoc. **(I)** Immunoblot images of MYBPC3 protein expression from human induced pluripotent stem cells (hiPSC), human induced pluripotent stem cell derived cardiomyocytes (hiPSC-CM), HeLa, HEK293T and NIH-3T3 cell lines. The total protein stain was used as the loading control. All results are shown as mean±SEM. Kruskal-Wallis test with Dunn’s post hoc test for multiple comparisons for was used for **B** and **H**. Brown-Forsythe and Welch ANOVA with Dunnett’s T3 multiple comparisons test was used for **D**. Unpaired t test with Welch’s correction was used for **F**.

We then wanted to determine if human and murine cell lines expressed MYBPC3 protein since it has been reported that some non-cardiomyocyte cell lines express sarcomere proteins.^12^ First, we compared human induced pluripotent stem cells (hiPSC) before and after differentiation into cardiomyocytes (hiPSC-CM). Undifferentiated hiPSC lacked detectable MYBPC3 or sarcomeric α-actinin protein, while hiPSC differentiated into cardiomyocytes had detectable MYBPC3 and sarcomeric α-actinin protein (**Figure 3I**). Non-cardiomyocyte cell lines from both humans and mice had no detectable MYBPC3 protein (**Figure 3I**). Overall, the results of these experiments show that cardiomyocytes are the cell source for MYBPC3 protein in the left ventricle.

### Cardiomyocyte versus germline *Mybpc3* deletion causes similar pathologic remodeling of the left ventricle

We then utilized our murine models to evaluate the impact of cardiomyocyte versus non-cardiomyocyte MYBPC3 protein expression on the development of cardiomyopathy. We found that left ventricular wall thickness and dilation were similar in mice with germline deletion of *Mybpc3* versus cardiomyocyte specific deletion of *Mybpc3* (**Figure 4A-C**). In addition, there were no significant differences detected between male and female mice (**Figure S3A-F**). Similar to the echocardiography results, heart mass (**Figure 4D**) and cardiomyocyte hypertrophy (**Figure 4E+F**) were similar between germline versus cardiomyocyte specific deletion of *Mybpc3*. In addition, we found similar levels of left ventricular fibrosis in mice with either germline or cardiomyocyte specific deletion of *Mybpc3* (**Figure 4G-H**). These results show that selective elimination of MYBPC3 protein in cardiomyocytes leads to similar pathologic remodeling of the left ventricle compared to elimination of MYBPC3 protein in all murine cells.

**Figure 4:**
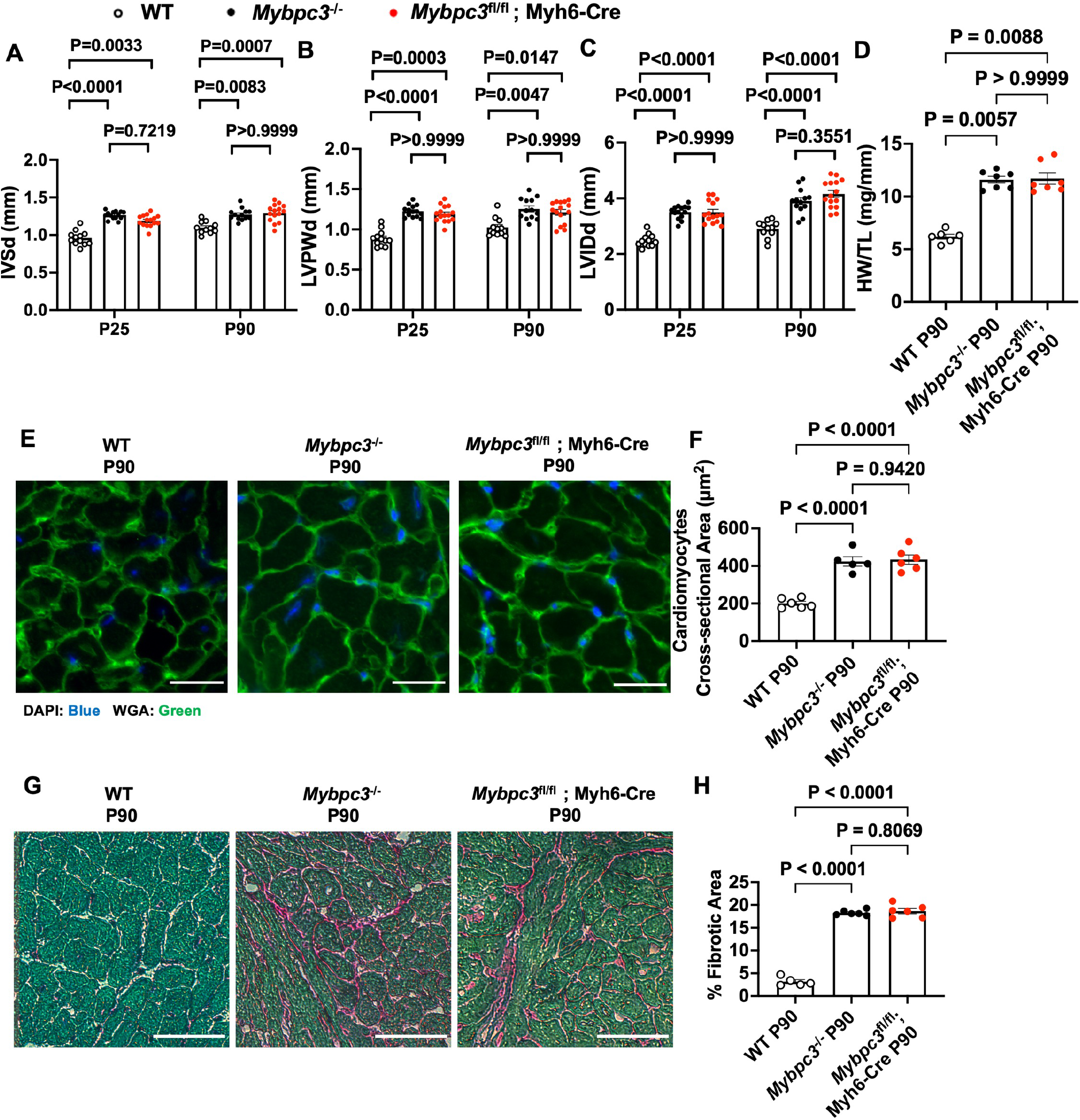
Cardiomyocyte versus germline *Mybpc3* deletion causes similar pathologic remodeling of the left ventricle. Transthoracic echocardiography was performed to measure **(A)** interventricular septal thickness at end-diastole (IVSd), **(B)** left ventricular posterior wall at end-diastole (LVPWd), **(C)** and left ventricular internal dimension at end-diastole (LVIDd) from WT (n=11-12), *Mybpc3*^−/−^ (n=13-14) and *Mybpc3*^fl/fl^; Myh6-Cre (n=15) at postnatal day 25 (P25) and P90. **(D)** Heart weight (HW) to tibia length (TL) ratio from WT (n=6), *Mybpc3*^−/−^ (n=7) and *Mybpc3*^fl/fl^; Myh6-Cre (n=7) at P90. **(E)** Representative images of wheat germ agglutinin (WGA) (green) and 4’6-diamidino-2-phenylindole (DAPI) (blue) fluorescence co-staining from WT (n=6), *Mybpc3*^−/−^ (n=5) and *Mybpc3*^fl/fl^; Myh6-Cre (n=6) at P90. Scale bars, 25μm. **(F)** Cardiomyocyte cross-sectional area quantification from WGA staining. Minimum of 100 cardiomyocytes/sample. **(G)** Representative images of Sirius Red/Fast Green staining from WT (n=5), *Mybpc3*^−/−^ (n=6) and *Mybpc3*^fl/fl^; Myh6-Cre (n=6) at P90. Scale bars, 25μm. **(H)** Myocardial fibrosis quantification from Sirius Red/Fast Green staining. All results are shown as mean±SEM. Kruskal-Wallis test with Dunn’s post hoc test for multiple comparisons was used for **A, B** and **D**. One-way ANOVA with Tukey’s multiple comparisons test was used for **F** and **H**. Brown-Forsythe and Welch ANOVA with Dunnett’s T3 multiple comparisons test was used for **C**.

### Cardiomyocyte versus germline *Mybpc3* deletion leads to similar abnormalities in left ventricular systolic and diastolic function

Next, we evaluated if cardiomyocyte specific versus germline deletion of *Mybpc3* led to differences in left ventricular function. We found that left ventricular systolic function decreased to a similar extent in both groups of mice in comparison to WT (**Figure 5A**). Likewise, left ventricular diastolic function was impaired to a similar degree in mice with cardiomyocyte *Mybpc3* deletion versus germline *Mybpc3* deletion (**Figure 5B-D**). Overall, these results show that elimination of MYBPC3 protein in cardiomyocytes versus all cell types leads to similar abnormalities in left ventricular systolic and diastolic function.

**Figure 5:**
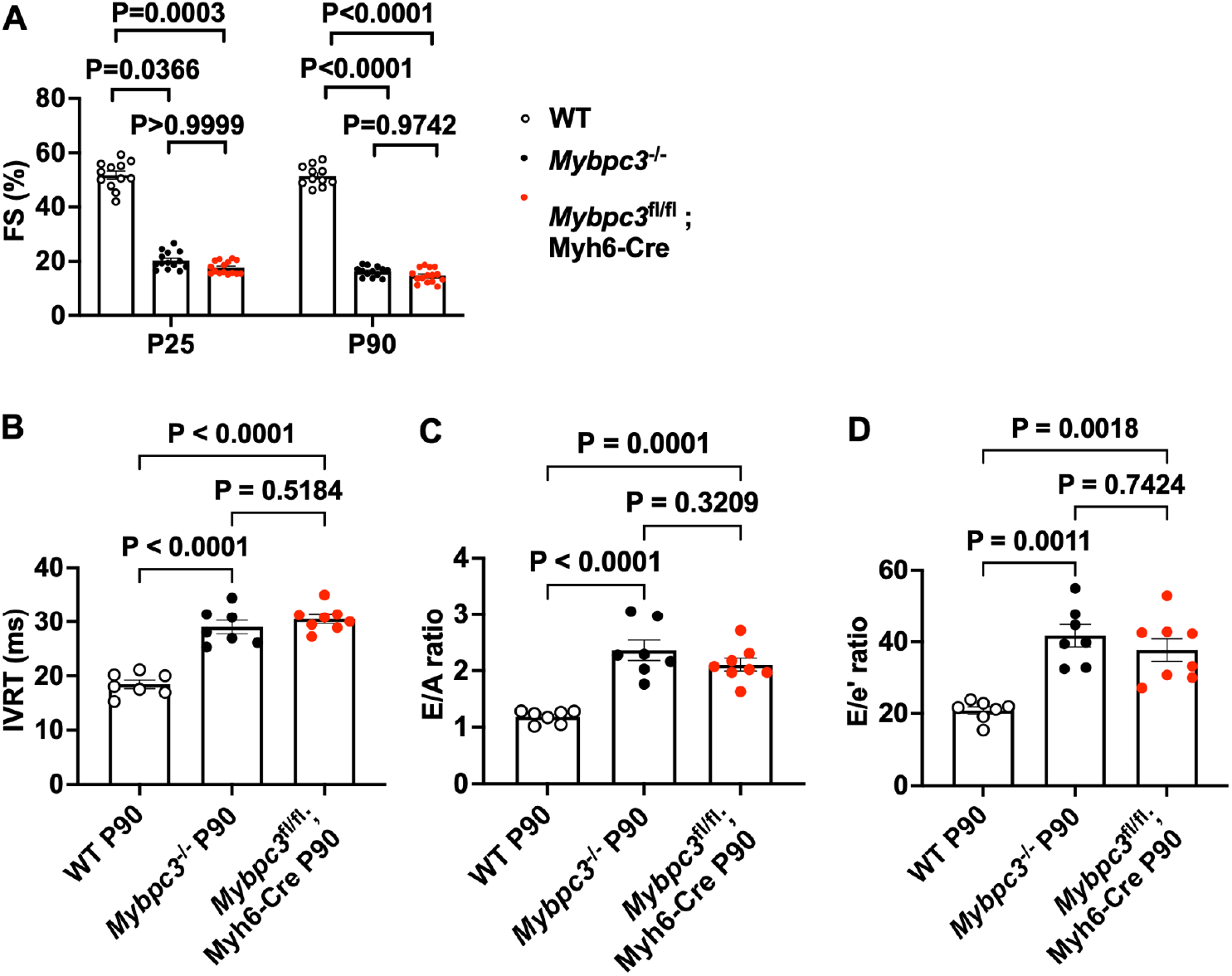
Cardiomyocyte versus germline *Mybpc3* deletion leads to similar abnormalities in left ventricular systolic and diastolic function. Transthoracic echocardiography was performed to measure **(A)** fractional shortening (FS) from WT (n=11-12), *Mybpc3*^−/−^ (n=13-14) and *Mybpc3*^fl/fl^; Myh6-Cre (n=15) at postnatal day 25 (P25) and P90. Transthoracic echocardiography was performed to measure **(B)** isovolumic relaxation time (IVRT), **(C)** mitral valve early to late filling velocity ratio (E/A) and **(D)** early transmitral valve flow velocity to early mitral annulus tissue velocity ratio (E/e’) from WT (n=7), *Mybpc3*^−/−^ (n=7) and *Mybpc3*^fl/fl^; Myh6-Cre (n=8) at P90. All results are shown as mean±SEM. Kruskal-Wallis test with Dunn’s post hoc test for multiple comparisons for was used for **A**. One-way ANOVA with Tukey’s multiple comparisons test was used for **B** and **C**. Brown-Forsythe and Welch ANOVA with Dunnett’s T3 multiple comparisons test was used for **D**.

## Discussion

Mutations in the sarcomere gene, *MYBPC3*, are one of the most common causes of hypertrophic cardiomyopathy in humans. Although MYBPC3 protein is an integral component of the sarcomere of cardiomyocytes, both *MYBPC3* mRNA and protein have been reported in non-cardiomyocyte cell populations.^10–12^ Therefore, we utilized transgenic murine models that enabled us to selectively eliminate cardiomyocyte *Mybpc3* mRNA. This allowed us to determine the cell source of extracardiac *Mybpc3* mRNA and investigate whether non-cardiomyocyte gene expression influenced the development of MYBPC3 related cardiomyopathy. We found that *Mybpc3* mRNA was present in multiple extracardiac organs, but cardiomyocytes were the primary source of this extracardiac *Mybpc3* mRNA. In addition, we found that despite the presence of extracardiac *Mybpc3* mRNA, there was no detectable MYBPC3 protein in extracardiac organs. Likewise, within the heart, there was no detectable MYBPC3 protein in non-cardiomyocyte cell populations. Importantly, cardiomyocyte specific elimination of MYBPC3 protein led to a similar effect on cardiac structure and function compared to elimination of MYBPC3 in all cell types. Taken together, this study shows that cardiomyocytes are the primary source of extracardiac *Mybpc3* mRNA but this extracardiac gene expression does not lead to detectable MYBPC3 protein or impact the development of cardiomyopathy related to MYBPC3 deficiency.

Extracardiac *MYBPC3* mRNA has been detected in whole blood samples in human patients with HCM and has been utilized to determine the impact of *MYBPC3* mutations on mRNA splicing.^17,18^ We found that both humans and mice had detectable *MYBPC3* mRNA in multiple extracardiac organs. However, it was unclear what the cell source was for this extracardiac *MYBPC3* mRNA. We utilized our transgenic murine models to determine that extracardiac *Mybpc3* mRNA is primarily derived from cardiomyocytes since the elimination of cardiomyocyte *Mybpc3* mRNA expression caused a significant decrease in extracardiac *Mybpc3* mRNA levels. This suggests that cardiomyocytes can secrete *Mybpc3* mRNA or passively release it through cardiomyocyte rupture. Interestingly, it was found that blood levels of *MYBPC3* RNA increased in patients after ST-elevation myocardial infarction suggesting that cardiomyocyte death may be one mechanism through which *MYBPC3* mRNA is released into the blood.^19^ Likewise, cardiomyocyte derived exosomes have been shown to contain both mRNA and DNA.^20^ The mechanisms controlling *MYBPC3* mRNA release and the biological role of circulating *MYBPC3* will need further investigation.

Interestingly, we detected residual low levels of *Mybpc3* mRNA in lung tissue after cardiomyocyte *Mybpc3* mRNA expression was eliminated. The cell source of this non-cardiomyocyte *Mybpc3* mRNA remains unclear but low levels of *MYBPC3* mRNA have been detected in human macrophages and neutrophils.^21,22^ Likewise, it was previously shown that Epstein-Barr virus immortalized lymphocytes can transcribe sarcomere genes such as myosin heavy chain 7 (MYH7).^23,24^ Importantly, despite detecting *Mybpc3* mRNA in lung tissue, we did not detect MYBPC3 protein in lung tissue using multiple different MYBPC3 antibodies.

It was previously reported that MYBPC3 protein can be detected in myocardial derived fibroblasts and NIH-3T3 cells.^12^ In our transgenic models that had cardiomyocyte specific deletion of *Mybpc3*, we were unable to detect residual MYBPC3 protein using in situ immunofluorescence, proximity ligation assays of myocardial tissue sections or immunoblotting of myocardial tissue lysate. Importantly, myocardial tissue samples from animals with cardiomyocyte deletion of *Mybpc3*, had similar levels of myocardial fibrosis compared to animals with germline deletion of *Mybpc3* in all cells. These results suggest that there were similar levels of pathologic fibroblast activation between the two models. Likewise, we were unable to detect MYBPC3 protein in NIH-3T3 cells cultured under standard culture conditions. Overall, our results show that non-cardiomyocyte cell populations of the murine myocardium do not readily express MYBPC3 protein.

We found that cardiomyocyte specific deletion of *Mybpc3* led to similar changes in cardiac structure and function compared to germline deletion of *Mybpc3* in all cells. This suggests that the small amount of residual *Mybpc3* mRNA expression in non-cardiomyocyte cell populations has no discernable impact on cardiomyopathy development and progression in preclinical murine models. These findings are particularly important because viral vector-based approaches under development to treat human cardiomyopathies related to MYBPC3 protein deficiency selectively target cardiomyocytes but not other cell populations.^15,16^

## Conclusion

Taken together, this study shows that cardiomyocytes are the primary source of extracardiac *Mybpc3* mRNA and MYBPC3 protein is localized to the cardiomyocyte cell population. Importantly, the expression of *Mybpc3* mRNA in non-cardiomyocyte cell populations has no discernible effect on the pathogenesis of cardiomyopathy resulting from MYBPC3 protein deficiency. Therefore, selectively increasing cardiomyocyte MYBPC3 protein levels should be the most efficient approach to mitigate the development of cardiomyopathies caused by MYBPC3 deficiency.

## Methods

### Ethics approval

All the mice that were used in this study were housed in animal facility accredited by the American Association for the Accreditation of Laboratory Animal Care (AAALAC). The animal experiments were conducted in accordance with the practices defined in the Guide for the Care and Use of Laboratory Animals which were approved and overseen by the University of Pittsburgh Institutional Animal Care and Use Committee (IACUC).

### Mouse models

To determine the impact of loss of MYBPC3 protein in all cells, we utilized a murine model that has a germline deletion of *Mybpc3* exons 3 to 5 (Mybpc3^−/−^).^25^ To achieve specific deletion of *Mybpc3* exons 3 to 5 in cardiomyocytes we crossed a Mybpc3^fl/fl^ line with the cardiomyocyte specific Myh6-Cre line (Jackson Labs, 011038). The generation of the Mybpc3^−/−^ and Mybpc3^fl/fl^ lines were previously described.^25^

### Human tissue analysis

Human control (unused donor) myocardial tissue samples were obtained in a deidentified manner from an institutional review board-approved tissue biorepository.

### Cell lines and culture

The human induced pluripotent stem cells (hiPSCs) were obtained from the Standford SCBI BioBank (SCVI274) and differentiated to hiPSC-derived cardiomyocytes (hiPSC-CMs) as previously described.^26^ Briefly, hiPSCs were maintained in essential 8 medium (A1517001, Gibco). Once cells were confluent (~80%), 10μM CHIR99021 (Selleckchem) was added in RPMI 1640 media with B27 minus insulin (Gibco) for 48 hours. The media was then changed to 5 μM IWR-1 (Sigma) in RPMI 1640 with B27 minus insulin for 48 hours. Cells were then maintained with RPMI 1640 with B27 minus insulin every other day until differentiated cells started to beat between 10 to 15 days post-differentiation. After 15 days of differentiation, iPSC-CMs were selected by RPMI 1640 without glucose supplemented with sodium L-lactate (Sigma) every other day until day 19 post-differentiation. The cells were then given RPMI 1640 with glucose and insulin transferrin selenite (ITS) plus media supplement (AR014; R&D systems) every other day. CMs were maintained with 5% CO_2_ at 37°C in a humidified incubator, and 30 days post-differentiated matured cells were used for the experiment. The NIH-3T3 (mouse embryonic fibroblast cell line), HeLa (human cervical carcinoma epithelial cell line) and HEK293T (human kidney epithelial cell line) cell lines were cultured separately in DMEM media (GIBCO) supplement with 10% fetal bovine serum (FBS) with 5% CO_2_ at 37°C in humidified incubator.

### Echocardiography

To assess the mouse cardiac systolic function and structure we performed transthoracic echocardiography using a Vevo 3100 (VisualSonics, Inc) without anesthesia. The interventricular septal thickness at end-diastole (IVSd), the left ventricular posterior wall at end-diastole (LVPWd), the left ventricular internal dimensions at end-systole (LVIDs) and at end-diastole (LVIDd) were acquired from M-mode short axis images. The left ventricular fractional shortening (FS) was obtained with the following formula (LVIDd – LVIDs)/LVIDd.

The mouse diastolic function was measured with B-mode long axis four chamber view under continuous isoflurane anesthesia. The left ventricular relaxation peak velocity in early diastole (E) and atrial contraction peak velocity in late diastole (A) was measured by mitral valve doppler flow. Early (E’) and late (A’) diastolic mitral annular tissue velocity was assessed by tissue Doppler image analysis. Finally, the Isovolumic relaxation time (IVRT) was obtained by measuring the period between the aortic valve closure and the mitral valve opening.

### Euthanasia and heart mass assessment

Mice were sedated with 5% isoflurane, and once unresponsive to toe pinch, body weight was measured. Before excising the heart, 500μL of blood was collected from the left ventricle (LV) or aortic puncture. The heart was then removed and heart weight was recorded. Mouse left ventricle tissue was isolated and then snap-frozen in liquid nitrogen for RNA and protein analysis. Full hearts were embedded in OCT for histology and immunohistochemistry staining. All the tissue samples were banked in −80°C freezer prior to being processed in future experiments. Mouse lower extremities were then amputated at the mid femur level and boiled to remove excess tissues and isolate the tibia. Digital calipers were used to measure the tibia length. Heart weight to tibia length (mm) ratios were then calculated.

### Histology

To determine cardiac fibrosis deposition, 5-μm thick sections were collected by cryo-sectioning the OCT-embedded heart. The harvested sections were then fixed for 5 minutes at −20°C in Acetone, then air-dried for 20 minutes at room temperature prior to being rehydrated in 1X PBS wash for 1 minute. Sections were then stained at room temperature for 1 hour with Sirius Red/Fast green solution (Chondrex 90461) before being washed in distilled water.

Sections were then imaged with a Zeiss Axioplan microscope to get bright-field image at x40 magnification. 10 pictures were taken per tissue sample. To determine the percentage of fibrotic area these pictures were analyzed with ImageJ software. To assess the fibrotic area, tissue stained with Sirius red was measured. Additionally, the myocardial area was determined by measuring the red and green stained tissue. The percentage of fibrotic area was then obtained by establishing a ratio of the fibrotic area per myocardial area for the tissue section of interest.

### Immunofluorescence and Wheat Germ Agglutinin Staining

Hearts were embedded in OCT prior to being sectioned at 5-μm on a cryostat (Thermo Fisher Scientific) and then harvested on Superfrost Plus Gold microscope slides (Fisher Scientific). Sections were fixed for 15 minutes with 4% paraformaldehyde then permeabilized with 0.2% Triton X-100 for 15 minutes with subsequent PBS washes. For the MYBPC3-WGA co-staining, an additional blocking step was performed by incubating the tissue sections with 0.1mg/mL Fab fragment goat anti-mouse IgG (Jackson Immuno Research, 115-07-003) at room temperature for 1 hour. Sections were then incubated for 1 hour with blocking buffer (1% BSA in 1X PBS). Sections were then incubated overnight at 4°C with MYBPC3 antibody (Santa Cruz Biotechnology, sc-137180, E-7). The following day tissue sections were washed and incubated for 1 hour at room temperature with fluorescent secondary antibody goat anti-mouse Alexa fluor 594 (Thermo Fisher Scientific, A-11032), goat anti-rabbit Alexa fluor 594 (Thermo Fisher Scientific, A-11005) and then washed in 1X PBS. Sections were then stained with wheat germ agglutinin (WGA) (Thermo Fisher Scientific, W6748) 1:200 at room temperature for 1 hour with subsequent washes in 1X PBS. Sections were then counterstained and mounted with Prolong Gold Antifade with 4’6-diamidino-2-phenylindole (DAPI). Slides were then imaged under wide-field fluorescent microscope (Zeiss) at x40 magnification. MYBPC3 fluorescence was determined by using the following equation: corrected MYBPC3 fluorescence = integrated density – (area of cardiomyocyte x mean of fluorescence of background readings), where integrated density is fluorescence intensity of the defined region of interest, area of cardiomyocyte is the size of the defined region of interest and mean of fluorescence background is the average intensity of 3 background regions of interest.

### Proximity ligation assay

In situ Proximity ligation assay (PLA) was performed as previously described.^27^ Mouse and human frozen heart tissue sections were cryosectioned 5μm thick. They were then fixed with 4% PFA for 15 minutes and washed with 1X PBS. The tissue sections were permeabilized with 0.2% Triton X-100 with subsequent PBS washes. Sections were then blocked using Duolink blocking buffer (Sigma) for 1 hour at RT and incubated with primary antibodies mouse MYBPC3 (Santa Cruz Biotechnology, sc-137180, E-7) 1:1000, and rabbit MYBPC3 (Invitrogen, 703574, 19H1L3) 1:1000 diluted in Duolink antibody diluent, overnight at 4°C. The sections were then incubated with secondary antibodies labeled with PLA probes (anti-mouse MINUS and anti-rabbit PLUS) diluted 1:5 in 1X Duolink Antibody Diluent buffer at 37°C for 1 hour. The ligation step was performed by adding ligase diluted in ligation buffer to the heart tissue section at 37°C for 30 minutes. The amplification step was then performed by adding polymerase diluted in amplification buffer in the dark at 37°C for 100 minutes. Secondary antibody incubation, ligation, and amplification steps were all completed in a humidity chamber. After final PBS washes, the heart tissue sections were incubated for 1 hour with blocking buffer (1% BSA in 1X PBS) and incubated with sarcomeric α-actinin antibody (Abcam, Ab9465) (1:1000) overnight at 4°C. After overnight incubation, the tissue sections were then washed and incubated for 1 hour at room temperature with fluorescent secondary antibody goat anti-mouse Alexa fluor 488 (Thermo Fisher Scientific, A-10667) followed by subsequent PBS washes. The heart tissue sections were then mounted with Prolong Gold Antifade with 4’6-diamidino-2-phenylindole (DAPI) (Thermo Fisher).

Slides were imaged under a wide-field fluorescent microscope (Zeiss) at 40x magnification. MYBPC3 complexes were estimated by thresholding the MYBPC3 PLA signal to measure the amount of MYBPC3 complexes located in the myocardium and the interstitial space. The average area, in pixels, of 25 individual PLA dots was measured across three randomly selected areas of the myocardium. In ImageJ, images were converted to RGB stack format. The percentage of red fluorescence in each image was thresholded using the ImageJ threshold function using an upper limit of 255, and a lower limit that was adjusted for each image to fully capture all of the red fluorescence. The percentage of red fluorescence was then multiplied by the total number of pixels in the image to calculate the total red pixels per image. This value was divided by the average PLA dot area to estimate the amount of MYBPC3 complexes per image. Non-cardiomyocyte MYBPC3 complexes were counted manually and subtracted from the total number of complexes to obtain the number of cardiomyocyte specific MYBPC3 complexes per image. The cardiomyocyte area in µm^2^ was measured by thresholding the sarcomeric α-actinin stained area (green channel) to obtain a percentage of green fluorescence in each image, which was then multiplied by the total image area. The non-cardiomyocyte area was calculated by subtracting the cardiomyocyte area from the total image area. Cardiomyocyte MYBPC3 complexes per 100 µm^2^ were determined by dividing the number of cardiomyocyte MYBPC3 complexes by the cardiomyocyte area (µm^2^) and then multiplying by 100. Non-cardiomyocyte MYBPC3 complexes per 100 µm^2^ were obtained by dividing the non-cardiomyocyte MYBPC3 complexes by the non-cardiomyocyte area and multiplying by 100.

### Human bulk tissue RNA-sequencing

*MYBPC3* read counts for heart left ventricle, whole blood, lung, liver, kidney cortex, brain frontal cortex and skeletal muscle were obtained from the GTEx Portal on 08/05/2024 and dbGaP accession number phs000424.v8.p2 on 08/05/2024. The *MYBPC3* RNA expression were normalized as a Log10(read counts).

### qRT-PCR

50mg of mouse left ventricle tissue was homogenized in 500μL of TRIzol®, while 500μL of mouse blood (harvested from LV and aorta puncture) were mixed with 500μL of TRIzol®. Then RNA extraction was performed following the protocol provided by the manufacturer (Direct-zol^TM^ RNA Miniprep, Zymo Research). Reverse transcription was performed following the manufacturer directions (Verso cDNA synthesis kit, Thermo Fisher Scientific). cDNA synthesis was performed using random hexamer primers except when specifically detailed as oligo dT primers. To assess gene expression, qPCR was performed with the Syber Green Master Mix (Applied Biosystem, A25742), Mybpc3 specific primers were designed to bind to the splice junctions of exons 2+3 and 3+4 (**Table S1**). cDNA samples were not diluted prior to running a 40 cycle qPCR reaction. Plate was set up using Quant-Studio-5 Real Time PCR System (Applied Biosystem). Each sample was run in duplicates or triplicates, and the Ct value was normalized using the housekeeping gene *Rpl32*. The 2^-ΔΔCt^ method was used to calculate the fold change in Mybpc3 mRNA expression relative to indicated group. Since the maximum cycle number was 40 for the qPCR, a Ct value of 40 was assigned to samples that were reported as undetected.

### Protein Electrophoresis and Western Blot

30mg of mouse left ventricle tissue were homogenized in 300μL ice-cold RIPA buffer (Sigma) supplemented with protease/phosphatase inhibitors (Thermo Fisher Scientific, 78441). 50μg of proteins from mouse left ventricle and 30μg of proteins from cell lines were separated on SDS-PAGE using 10% Tris-Glycine gel (Bio-Rad, 4561093, 5671033) and transferred to a 0.45-μm low-fluorescence PVDF membrane (Bio-Rad) at 4°C for 1 hour at 100V. Total protein was imaged on the membrane with the Revert^TM^ 700 total protein stain kit (LI-COR, 926-11016). Membrane was then blocked with 5% BSA or non-fat dry milk in TBS/Tween20 (0.1% v/v) at room temperature for 1 hour. Primary antibody was incubated at 4°C overnight with MYBPC3 (Santa Cruz Biotechnology, sc-137180, E-7), (Invitrogen, 703574, 19H1L3), or β-actin (Cell Signaling Technology, 8457) in 1:500. The following day membranes were washed with TBST and incubated with secondary antibody goat anti-mouse HRP (Cell Signaling Technology, 7076S), or goat anti-rabbit HRP (Cell Signaling Technology, 7074S) at room temperature for 1 hour with subsequent washes and then imaged with Chemi Doc apparatus (Bio-Rad) using Clarity ECL Substrate (Bio-Rad, 1705061). For LI-COR imaging system, goat anti-mouse (IRDye 680 LT, 926-68020) and goat anti-rabbit (IRDye 800 CW, 926-32211) were used, and membranes were imaged with Odyssey CLx imaging system (LI-COR).

### Statistical analysis

All experimental data are displayed as mean ± SEM. The normal (Gaussian) distribution of the experimental data set was tested with the Shapiro-Wilk normality test. If the data set were normally distributed, statistical significance between two experimental groups was tested using two-tailed unpaired Student’s t-test. However, if the data set failed to pass the F test to compare variances, therefore, a Welch’s t-test was performed. For the data set including more than two experimental groups a one-way ANOVA with a Tukey’s multiple comparisons test was performed to test the statistical significance. However, if the data set failed to pass the F test to compare variances, therefore, a Brown-Forsythe and Welch ANOVA with Dunnetts’s T3 multiple group comparisons test was performed. For data sets that were not normally distributed, Mann-Whitney U test was used to test statistical significance between two experimental groups. However, Kruskal-Wallis test with Dunn’s post hoc test for multiple group comparison for data sets including more than two experimental groups. All statistical analysis was performed with Prism 10 software (GraphPad). Statistical significance was considered a p-value less than 0.05.

## Non-standard Abbreviations and Acronyms

HCM: hypertrophic cardiomyopathy
LV: left ventricle
LVH: left ventricular hypertrophy
MYBPC3: myosin binding protein C3

## Acknowledgements

We would like to acknowledge Dr. Sruti Shiva (Heart, Lung, Blood Vascular Institute, University of Pittsburgh) and Dr. Yael Nechemia-Arbely (Hillman cancer center, University of Pittsburgh) for providing us with NIH-3T3 and HeLa cells respectively.

## Author contributions

N.G.C and J.R.B designed the research study. N.G.C., K.P.L., S.T.M, J.H.K., K.S., S.P., and J.R.B. conducted experiments and data analysis. N.G.C., K.P.L., S.T.M, J.H.K., K.S., S.P., and J.R.B. prepared and edited the manuscript.

## Source of Funding

This work was supported by grants from the National Institutes of Health (HL136824, HL160890, HL167955, and HL169784 to J.R.B.)

## Disclosures

None.

## Supplementary Data

Figure S1, S2, S3 and Table S1

## Supplementary Figure Legend

**Figure S1.**
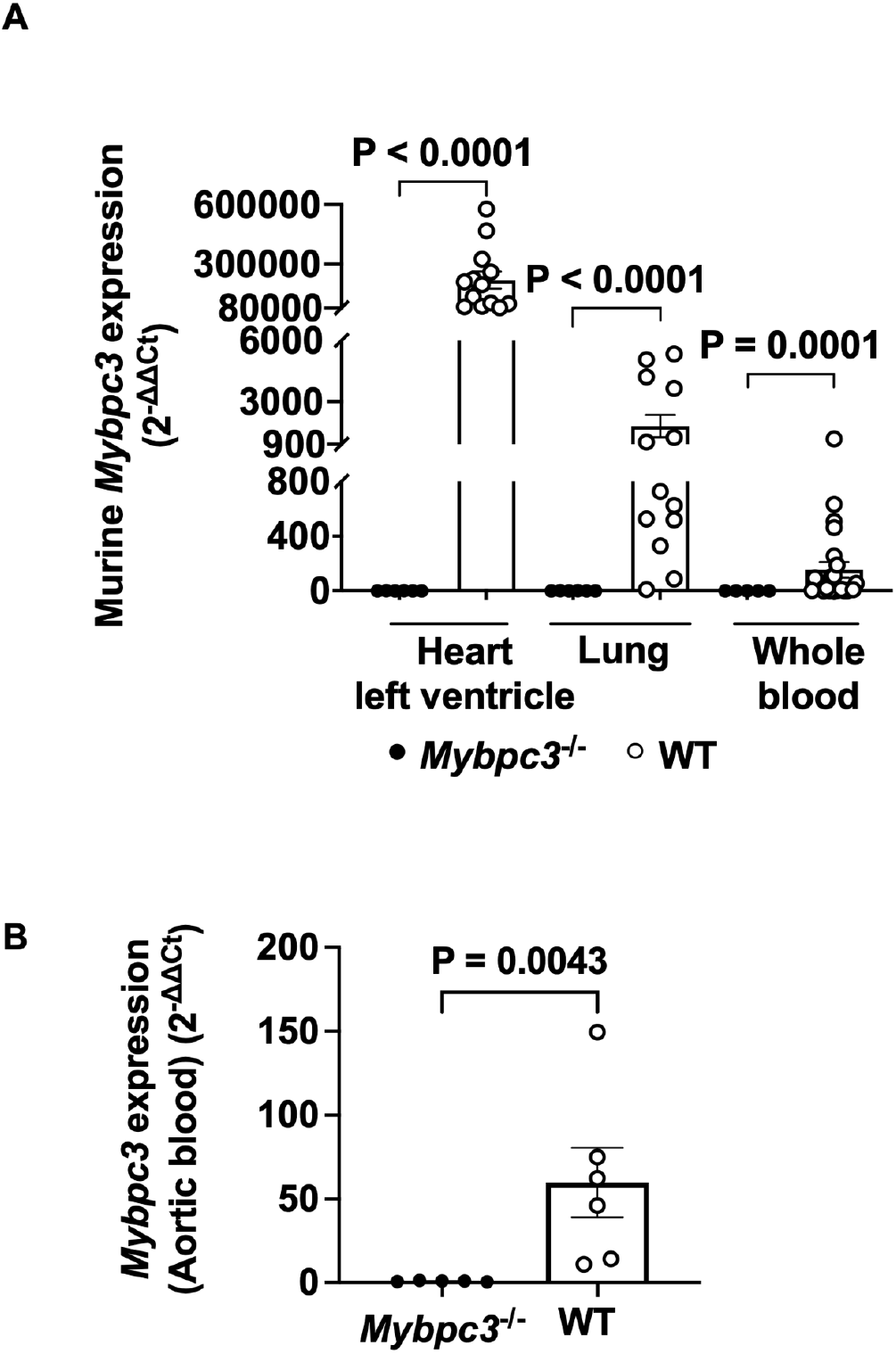
(**A**) Quantification of *Mybpc3* expression from *Mybpc3*^−/−^ and WT heart left ventricle (n=6-13), lung (n=6-13) and whole blood (n=5-24) at P90. The *Mybpc3* gene expression was normalized to the housekeeping gene *Rpl32* and the cDNA was created using oligo(dT) primers. **(B)** Quantification of *Mybpc3* mRNA expression from *Mybpc3*^−/−^ and WT whole blood from aortic puncture (n=5-6), at postnatal day 90 (P90). The *Mybpc3* gene expression was normalized to the housekeeping gene *Rpl32*. Results are expressed as a fold change relative to *Mybpc3* mRNA expression in the *Mybpc3*^−/−^ mouse tissue. All results are shown as mean±SEM. Mann-Whitney U test was utilized for **A + B**.

**Figure S2.**
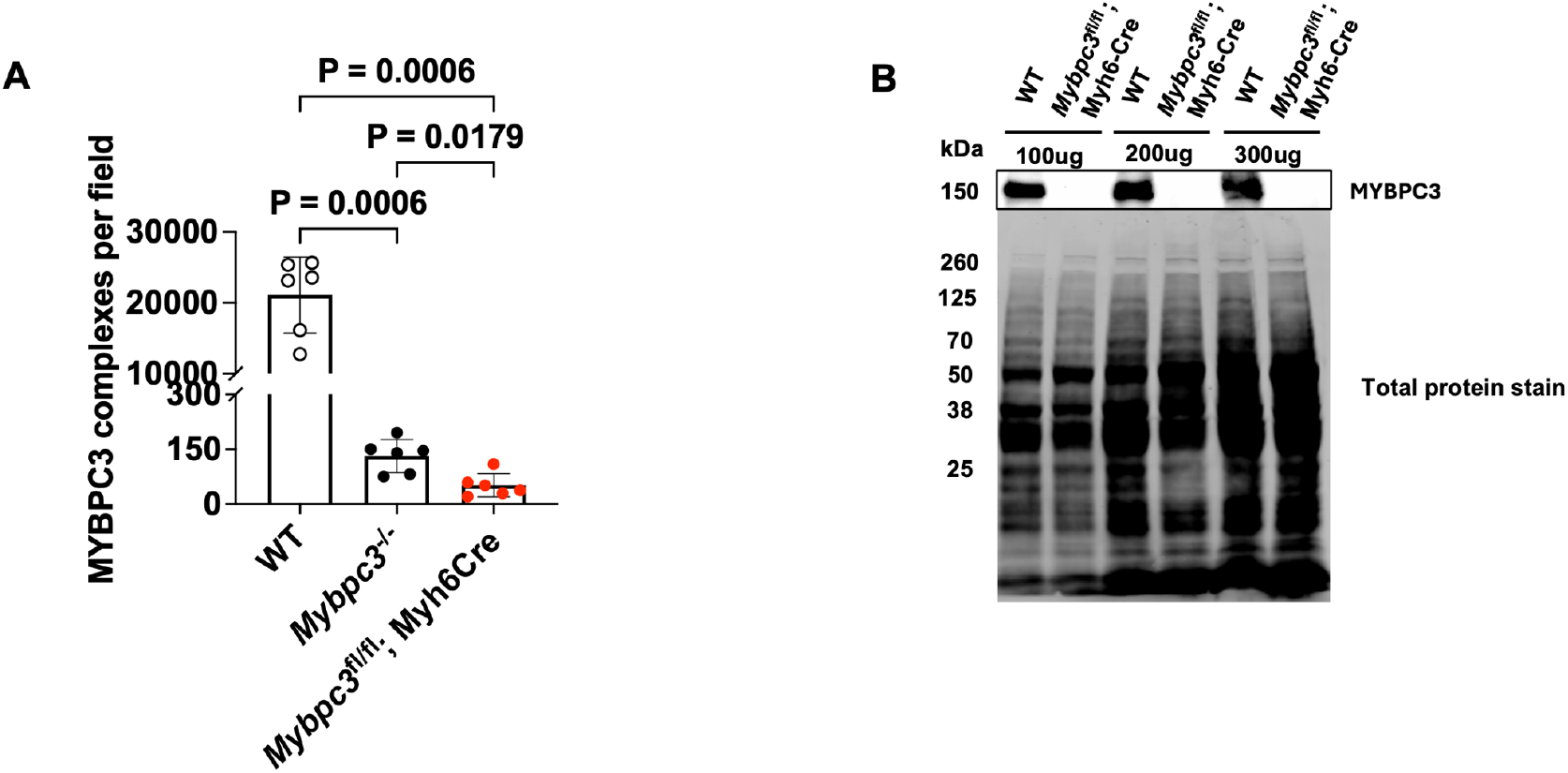
(**A**) In situ proximity ligation assay (PLA) quantification for MYBPC3 complexes per field in left ventricular tissue from WT (n=6), *Mybpc3*^−/−^ (n=6), and *Mybpc3*^fl/fl^;Myh6Cre (n=6) at P180. **(B)** Immunoblot images of MYBPC3 protein from WT and *Mybpc3*^fl/fl^; Myh6-Cre left ventricular tissue using different concentrations of total protein lysate (100μg, 200μg and 300μg). The total protein stain was used as the loading control. All results are shown as mean±SEM. Brown-Forsythe and Welch ANOVA with Dunnett’s T3 multiple comparisons test was used for **A**.

**Figure S3.**
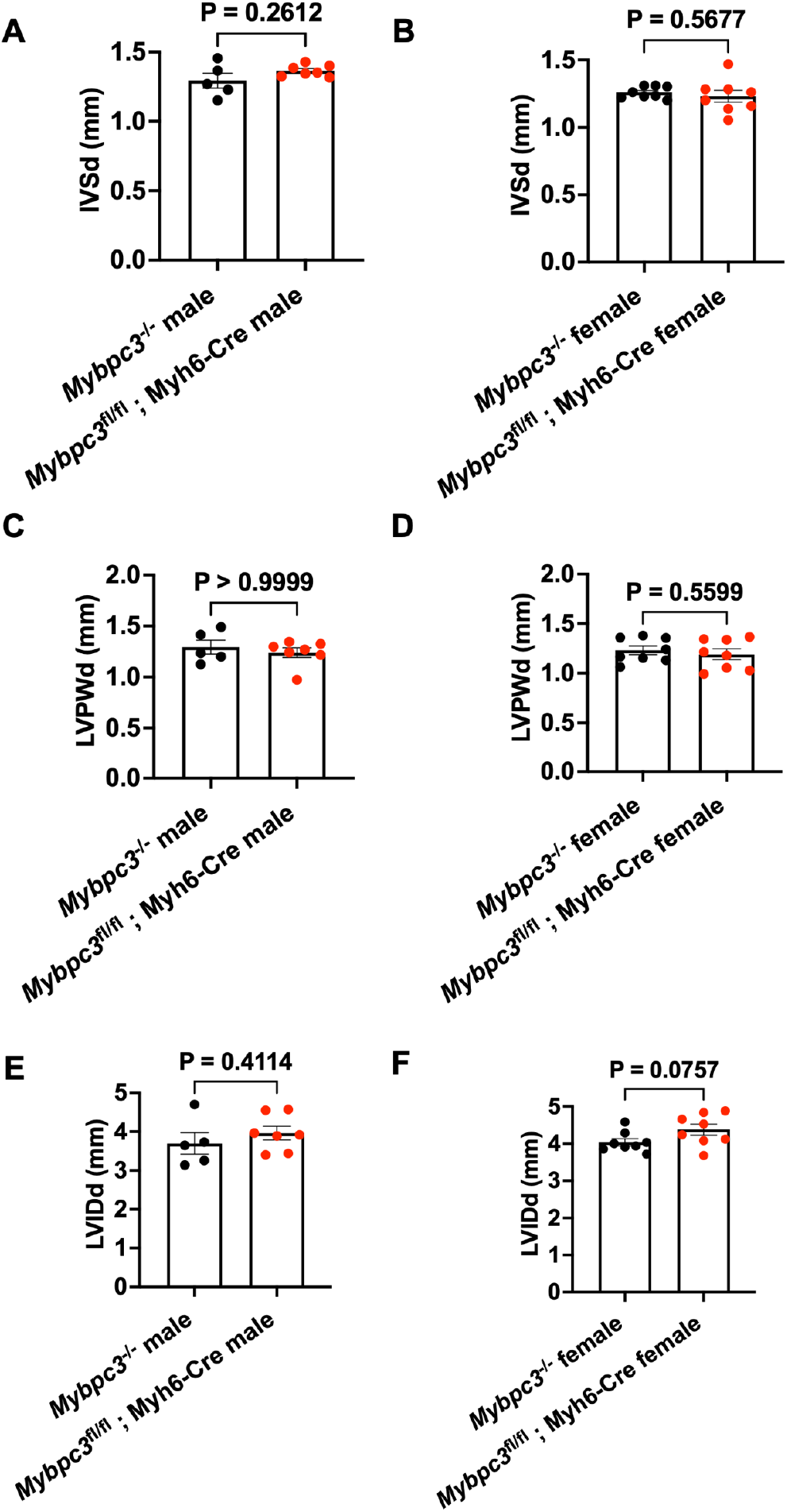
Transthoracic echocardiography was performed to measure **(A)** interventricular septal thickness at end-diastole (IVSd), **(B)** left ventricular posterior wall at end-diastole (LVPWd), **(C)** and left ventricular internal dimensions at end-diastole (LVIDd) from *Mybpc3*^−/−^ (male n=5; female n=8) and *Mybpc3*^fl/fl^; Myh6-Cre (male n=7; female n=8) at P90. All results are shown as mean±SEM. Student’s Welch’s t test was used for **A** and **B**. Mann Whitney U test was for **C**. Unpaired Student’ t test was used for **D, E** and **F. Supplementary Table Legend**

**Table S1.**
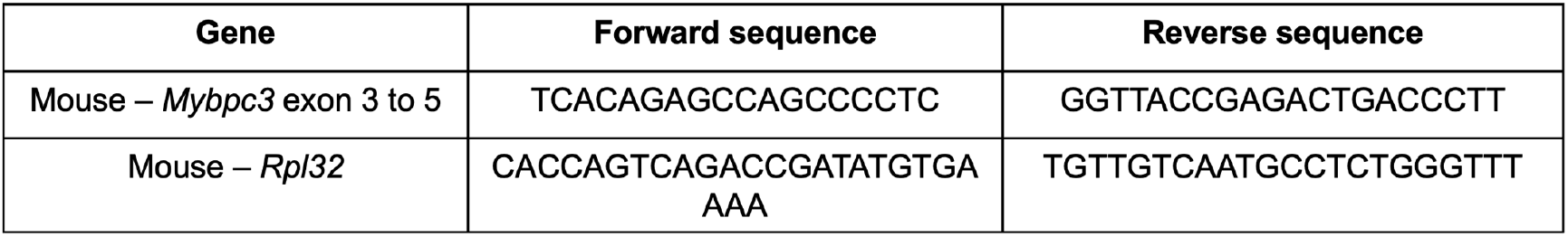
List of oligonucleotide primer sequences used for qRT-PCR.

## Notes

### Competing Interest Statement

The authors have declared no competing interest.

